# Guiding a language-model based protein design method towards MHC Class-I immune-visibility profiles for vaccines and therapeutics

**DOI:** 10.1101/2023.07.10.548300

**Authors:** Hans-Christof Gasser, Diego Oyarzun, Ajitha Rajan, Javier Alfaro

## Abstract

Proteins have an arsenal of medical applications that include disrupting protein interactions, acting as potent vaccines, and replacing genetically deficient proteins. While therapeutics must avoid triggering unwanted immune-responses, vaccines should support a robust immune-reaction targeting a broad range of pathogen variants. Therefore, computational methods modifying proteins’ immunogenicity without disrupting function are needed. While many components of the immune-system can be involved in a reaction, we focus on Cytotoxic T-lymphocytes (CTLs). These target short peptides presented via the MHC Class I (MHC-I) pathway. To explore the limits of modifying the visibility of those peptides to CTLs within the distribution of naturally occurring sequences, we developed a novel machine learning technique, `CAPE-XVAE`. It combines a language model with reinforcement learning to modify a protein’s immune-visibility. Our results show that `CAPE-XVAE` effectively modifies the visibility of the HIV Nef protein to CTLs. We contrast `CAPE-XVAE` to `CAPE-Packer`, a physics-based method we also developed. Compared to `CAPE-Packer`, the machine learning approach suggests sequences that draw upon local sequence similarities in the training set. This is beneficial for vaccine development, where the sequence should be representative of the real viral population. Additionally, the language model approach holds promise for preserving both known and unknown functional constraints, which is essential for the immune-modulation of therapeutic proteins. In contrast, `CAPE-Packer`, emphasizes preserving the protein’s overall fold and can reach greater extremes of immune-visibility, but falls short of capturing the sequence diversity of viral variants available to learn from. Source code: https://github.com/hcgasser/CAPE (Tag: `CAPE 1.1`)

## 1 Introduction

The recent SARS-CoV-2 pandemic has shown that RNA vaccines are safe, quick to develop and can reliably induce the production of viral proteins by patients. This success has spurred development of more vaccines against other pathogens [1]. Additionally, similar technologies could also be used to stimulate the production of therapeutic proteins exhibiting a broad range of functions. As of May 4, 2023, the pharmaceutical company Moderna is advancing mRNA therapeutics targeting conditions including autoimmune hepatitis, cancer, and genetic disorders like propionic acidemia and cystic fibrosis [1].

The desired interaction of these pharmaceutical proteins with the immune-system is application dependent. On one hand, an unintended immune-response to a therapeutic protein may undermine its efficacy or, in extreme cases, provoke autoimmune-reactions. Consequently, deimmunization of proteins has garnered attention in this context [2, 3, 4]. On the other hand, an immune-response is intended to the SARS-CoV-2 spike protein vaccine - ideally directed against a broad spectrum of viral variants. Taken together, there is a pressing need for techniques capable of integrating diverse immune-response objectives into the computational protein design process.

Research has so far mostly focused on removing antibody (Ab) surface epitopes [5] and decreasing the presentation of MHC Class II (MHC-II) epitopes crucial for B-cell activation [6]. However, gene therapies as well as mRNA therapeutics and vaccines translate into proteins within cells. From within the cell these proteins are presented to the immune-system via the MHC-I pathway, and thereby potentially triggering a CTL reaction. Therefore versatile, computational methods are also needed to modify the CTL reaction. This need is demonstrated by anti-transgene immunity [7, 8] and examples where vector capsid proteins flag hepatocytes for CTL destruction [9, 10] as well as examples of unwanted immune-responses to the bacterial-derived CRISPR-Cas9 gene editing machinery [11]. Conversely, vaccines - particularly those targeting rapidly mutating viruses with highly diverse quasi-species like HIV [12] - need to include sequences that expose the immune-system to a wide array of MHC-I immune-visible peptides, mirroring the diversity found in the viral population. This is underscored by Borrow et al. [13] who found evidence for rapid intra-patient evolution of HIV to evade CTL recognition leading to the conclusion that a key aim of HIV-vaccines should be to elicit strong CTL responses against multiple codominant viral epitopes. Nevertheless, immune-reaction to an antigen is a complex process and many aspects are not fully understood yet. Therefore, methods need to be flexible to incorporate the latest developments.

This article introduces the first stage of developing the Controlled Amplitude of Present Epitopes (CAPE) framework, which is focused on managing MHC-I visibility. In this article we make the following novel contributions. (1) We develop `CAPE-XVAE`, a flexible and novel machine learning (ML)-based approach that combines the abilities of Variational Autoencoders (VAEs) and reinforcement learning (RL), to generate MHC-I immune-visibility modified protein sequences. Its flexibility will allow extension to other components of the immune-system in the future. (2) We introduce `CAPE-Packer`, a modification of an existing physics-based approach from Yachnin et al. [6], enabling it to reduce or increase MHC-I immune-visibility. And finally, (3) we apply these two novel methods to the HIV Nef protein, comparing their ability to modify immune-visibility while maintaining stability and similarity to naturally occurring sequences. We find that `CAPE-XVAE` effectively modifies the immune-visibility of generated Nef proteins, while maintaining sequence similarity to natural proteins. In contrast `CAPE-Packer` achieves even greater immune-visibility modification, albeit it generates proteins markedly different from naturally occurring ones in sequence space.

### Box 1

**Terms and concepts**

- **Support sequences and natural sequences:** Often applying an analysis to all sequences in the dataset would be prohibitively computationally expensive. To still be able to describe the behavior of all sequences in the dataset, we once randomly selected `300` sequences from the whole dataset and apply the analysis just to those. We refer to these `300` sequences as *support sequences*, while we use *natural sequences* to refer to all sequences in the whole dataset.
- **Generated/designed sequences:** In contrast to natural sequences, we use generated- and designed sequences in a synonymous way to refer to those that were generated by a computational model.
- **Natural vs artificial peptides:** Natural peptides can be found in at least one natural sequence. Artificial ones cannot.
- **Closest support sequence:** For a given (query) sequence, *closest support sequence* refers to the support (subject) sequence with the highest pairwise TMscore. If the query sequence is a support sequence, it is excluded from the potential subject sequences.

### Box 2

**Terms and concepts - immune-visibility**

- **MHC-I immune-visibility (also just ‘immune-visibility’ or ‘visibility’):** The human immune-system is comprised of many components. Our study focuses on the MHC-I pathway which is used by CTL to detect destruction targets (see 2.1.1). We define the visibility of a protein to be the number of different 8, 9 and 10-mers within a protein sequence, that are predicted to be presented on the cell’s surface. Overlapping k-mers are counted separately. Also, if a k-mer is presented by several MHC-I alleles, then each one is independently increasing the visibility number by one. For this study we selected the following common MHC-I alleles to tailor it towards a hypothetical patient: HLA-A*02:01, HLA-A*24:02, HLA-B*07:02, HLA-B*39:01, HLA-C*07:01, HLA-C*16:01. Hence, a sequence’s visibility is the same as the sum of peptide rewards (see Subsection 2.5) under the *increased* visibility profile (see inside box below) over all 8 to 10 amino acid long peptides within the sequence. Importantly, while being visible/presented on the cell’s surface is a necessary condition for a peptide to cause an immune-reaction by CTLs, this is by no means a sufficient one (see 2.1.1).
- **Visibility profiles - *baseline* /*reduced* /*increased* /*inc-nat* :** We are trying to modify how the protein is visible to the immune-system. We use the word *baseline* to refer to generated sequences that were produced by a model without any immune-visibility modification. For `CAPE-XVAE`, this means just using the VAE [14] decoder without any modification step. For `CAPE-Packer` this means using an energy term that does not consider the visibility of the designed sequence. In contrast, we use *reduced/increased* to refer to generated sequences with (lower/higher) modified immune-visibility. In *inc-nat* (*increased-natural*) we modified the protein’s visibility profile by maximizing naturally occurring visible peptides and minimizing artificial ones. (see also Subsection 2.5, `Box1`)
- **Highly/lowly immune-visible sequences:** All natural or designed sequences that have an immune-visibility above/below the 99.5%/0.5% percentile of immune-visibility in the natural sequences.

## 2 Material and methods

This section introduces the methods and dataset employed in this paper, with essential terms summarized in `Boxes 1 and 2`. The first subsection delves into the MHC-I pathway. Following that, we present an approximation method designed to accelerate the prediction of peptides presented through this pathway. We then introduce the dataset used in our analysis. Subsequently, we shed light on the inner workings of our ML approach, `CAPE-XVAE`, and our physics-based approach, `CAPE-Packer`. Finally, we present a comprehensive overview of our evaluation process.

### 2.1 The MHC-I pathway and predicting its presented peptides

The MHC-I pathway is key in immune-response, presenting cellular peptides to T cells. Detailed exploration of this pathway and its broader immune-system integration are reviewed elsewhere, beyond the scope of this paper. Nevertheless, we provide a concise overview and refer the interested reader to Murphy and Weaver [15] for more in-depth information before introducing how we assessed peptide presentation via this pathway.

#### 2.1.1 The MHC-I pathway

Within most human cells ribosomes constantly produce proteins, while proteasomes constantly degrade them into small fragments, known as peptides. Some of these get transferred into the cell’s endoplasmic reticulum, where some bind to MHC-I proteins (typically 8 to 10 amino acids (AAs) long [15, Chapter 4-15]), forming pMHC complexes. These are transported and displayed on the cell’s surface for surveillance by CTLs.

CTLs use a T-cell receptor (TCR) to bind to these visible/presented pMHC. Due to random genetic recombination, the TCRs within a person’s population of CTLs vary - leading to differences in the specific pMHC they bind to. To prevent auto-immunity, TCR undergo selection in the thymus to be tolerant of the person’s own naturally occurring peptides while still binding specifically to foreign peptides. These foreign peptides can originate from viral proteins, cancer proteins, or other intra-cellular pathogens. Additionally, CTLs develop from naive CD8 T cells via priming. These T cells circulate the bloodstream visiting lymph nodes, where they transiently bind to antigen-presenting cells (typically dendritic cells). If a naive CD8 T cell encounters its cognate antigenic peptide presented on an antigen-presenting cell and receives co-stimulation [15, Fig. 9.22], the T cell proliferates and its progeny differentiate into CTLs. If a CTL detects a foreign peptide on a cell’s surface, it initializes the destruction of this cell. [15, Chapter 9] MHC-I alleles exhibit variation in peptide specificity. To ensure the presentation of a broad range of peptides on the cell surface and make them visible to CTLs, evolution has led to the expression of a diverse set of MHC-I proteins across the human population. In fact, three main genes (HLA-A, HLA-B, and HLA-C) are present in humans, allowing each individual to express up to six different MHC-I proteins [15, Chapter 6].

From the above, it becomes evident that *visibility* to CTL of a peptide is a necessary but not a sufficient condition for an immune-reaction. For example, presented self-peptides, as well as foreign peptides for which no matching TCR exists, will not trigger an immune-reaction. Additionally, an immune-reaction requires prior activation of the CTL. Therefore, *visibility* and immune-reaction are not the same.

#### 2.1.2 Assessing antigen presentation via MHC Class I

For the estimation of MHC-I immune-visibility, our study employs *netMHCpan-4*.*1* [16]. This tool represents the latest iteration in a series of MHC-I prediction models dating back to the early 2000s. Based on a given peptide sequence and a specific MHC-I allele, the model produces a score. Higher values indicate greater confidence in the peptide’s presentation by the MHC-I protein. The score’s reversed rank relative to random peptides, known as *EL-rank*, serves us to assess MHC-I presentation of the peptide. Peptides with an *EL-rank* of below 2% are considered “weak binding” and those below 0.5% are deemed “strong binding” ^1^.

In our exploration of methods to integrate the immune-visibility constraint into the protein design process, we encountered challenges with an approach suggested by Yachnin et al. [6]. This method involves numerous peptide presentation prediction requests, rendering direct use of *netMHCpan* in `CAPE-Packer` impractical (see Subsection 2.4). To reduce the execution time of the immune modification process in `CAPE-Packer` and `CAPE-XVAE` (while still relying directly on *netMHCpan* during evaluation), we approximated the behavior of *netMHCpan* using a position weight matrix (PWM) based approach.

To construct the PWMs, we initially generated a uniform sample of three million random peptides of lengths 8, 9, and 10 AAs. Subsequently, we employed *netMHCpan* to calculate the *EL-rank* for all considered MHC-I alleles, selecting peptides with an *EL-rank* below 2%. For each position within these peptides, we computed the distribution of AAs, obtaining a PWM for each MHC-I allele and peptide length. Next, we scored all the random peptides using these PWMs. This involved summing up the logarithmic probabilities assigned by the PWM to each AA at their corresponding positions within the peptide. The scores were then used to establish rankings for peptides, equivalent to the *EL-rank*.

### 2.2 Protein dataset

We applied our protein design methods to the HIV Nef protein. HIV’s urgent need for effective vaccines as well as Nef’s relatively short sequence length, slightly above 200 AAs, and the availability of approximately 25 thousand different sequence examples make it a suitable test case. On 6 June 2023, we retrieved unaligned HIV Nef sequences from the *Los Alamos National Laboratory* HIV Sequence Database ^2^. To avoid using dysfunctional sequences or sequences with sequencing errors, we only considered sequences between 200 and 220 AAs length. Many sequences occur multiple times and we assigned proportionally higher weights during `CAPE-XVAE` training to those. While ‘X’-containing sequences (unknown AA, absent in *complete sequences*) were retained for training, they were excluded from the evaluation process. We randomly partitioned our dataset into the following subsets:

### 2.3 CAPE-XVAE

The capabilities of VAEs and RL techniques are combined within `CAPE-XVAE` to sample proteins with a modified immune-visibility profile. The VAE‘s decoder is trained to generate meaningful examples for likely points of the latent space distribution. However, as seen in [17], just randomly sampling from this latent space distribution to generate protein sequences will typically only lead to marginal modifications in immune-visibility, as the immune-visibility of the generated proteins will mirror that of the training data.

Therefore, more efficient ways to navigate the latent space are necessary. For this purpose, our approach relays immune-visibility information through the decoder. Since back-propagation through the decoder’s discrete output is infeasible, we formulate our task as a RL problem. Unlike in standard RL, we do not modify the model parameters. Instead, we update the latent representation of a protein to better align with our desired immune-visibility profile (see `Box2`). The improved representation can then be converted to a sequence by the decoder. As mentioned earlier, VAEs’ decoders are trained to assign meaningful examples (protein sequences) to all likely points of the latent space distribution. Without restricting the latent representation updates, we observed that the RL algorithm would move them far away from the center of the distribution to very unlikely areas. Therefore, we restrict the updates to remain within a box around an initially sampled latent representation. This significantly improves the similarity of the generated sequences to natural ones, while still allowing for sufficient room for exploration. Since this still does not guarantee the generation of a meaningful protein in every instance, we employ an array of bioinformatics methods to filter out potentially errant sequences (see Subsection 2.6).

Similar to Dauparas et al. [18] we train the decoder to generate sequences in any ordering - allowing certain parts of the sequence to be fixed upfront. This, as well as the ability to adjust the modification procedure to a wide range of immune-visibility profiles, makes `CAPE-XVAE` a particularly flexible approach for protein design.

After presenting `CAPE-XVAE`’s architecture, we delve into its VAE training process. Finally, the immune-visibility modification procedure is outlined.

#### 2.3.1 Architecture

`CAPE-XVAE` is based on the VAE architecture [14] and, therefore, comprises the three main components listed below: an encoder, a latent space, and a decoder (see **Figure 1a**). The encoder and the decoder were realized as transformers [19]. More details on the various sub-components can be found in the Supplement **S.1.2**.

**Figure 1:**
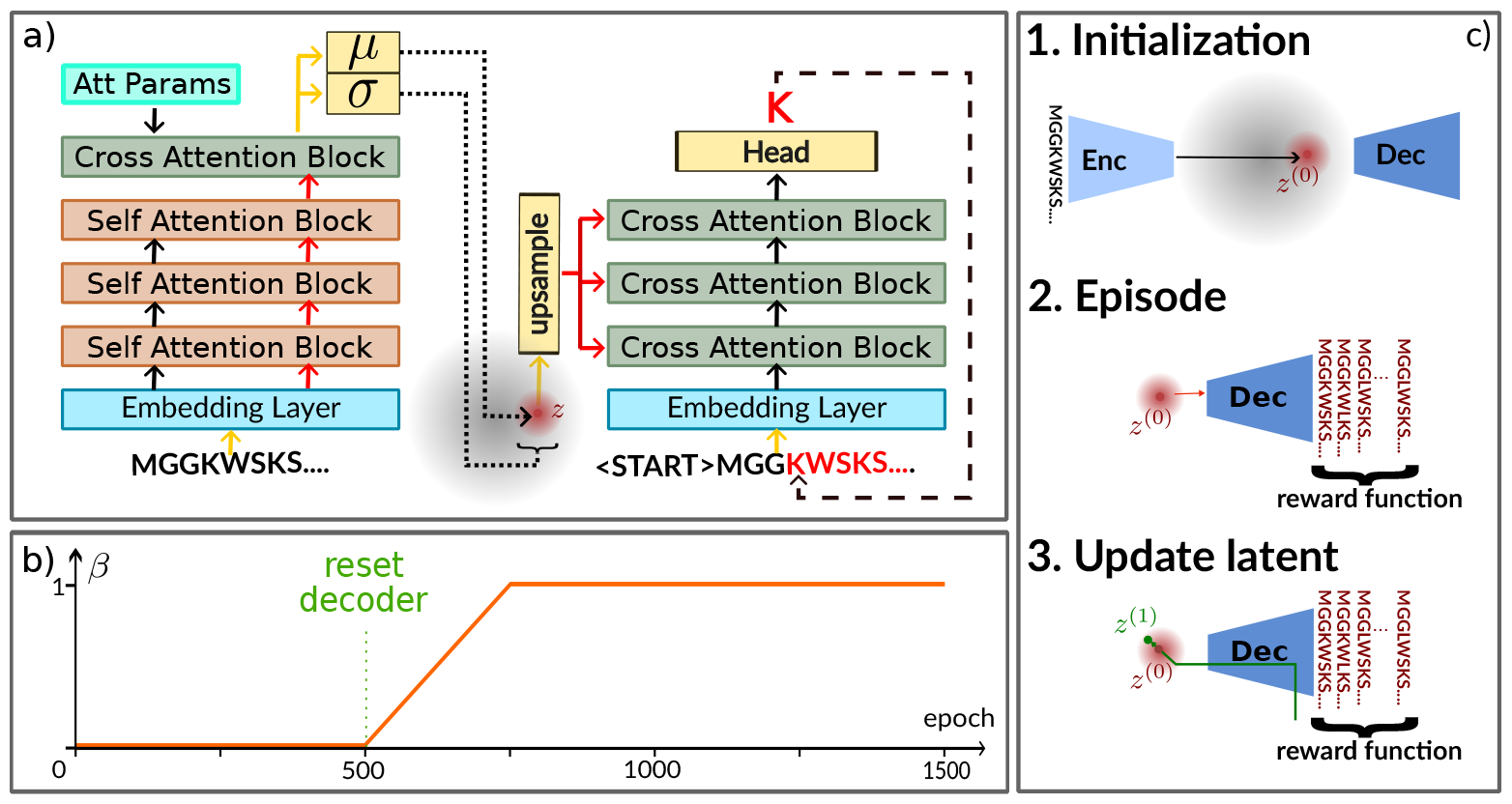
Overview of the CAPE-XVAE model architecture, training and immune-visibility modification process. (a) Sketch of the `CAPE-XVAE` architecture. On the left side is the encoder, in the middle the latent space and on the right side the decoder. More details can be found in the Supplement **S.1**. (b) Shows the evolution of the KL-divergence term weight during `CAPE-XVAE` training. (c) Depicts the steps followed during the immune-visibility modification process

- **Encoder:** `CAPE-XVAE`’s encoder consists of an *Embedding Layer*, an *Encoder Stack*, a *Cross Attention Block* and two linear transformations embedding the stack’s output into latent space. Along with a learnt positional encoding, each AA in the input protein sequence *x* is first encoded as a *d* dimensional vector using a learnt embedding layer. The *Encoder Stack* (comprising multiple *Self Attention Blocks*) processes these into a same length sequence of contextually encoded vectors. To create an information bottleneck, first a *Cross Attention Block* converts this sequence into *L < N* (currently *L* = 1) vectors (each of dimension *d*). Subsequently, two linear transformations further compress each vector to *l < d* dimensions. The first transformation *µ* generates a mean vector for the latent representation’s conditional distribution *p*(*·*|*x*), while the second one log(*σ*^2^) produces its log variance vector.
- **Latent Space:** To generate sequences, first a vector *z* needs to be sampled in latent space. Sampling from the latent’s conditional distribution is appropriate when *z* should represent a sequence similar to *x* (e.g. during training). In contrast, when we are only interested in generating any sequence similar to the training set, sampling from the unconditional distribution is possible. For the loss function, we assume this to be a standard normal distribution. However, we acknowledge that this assumption may not accurately represent the distribution of natural sequences in latent space. So, when generating new *baseline* (see `Box2`) examples, we calculate the mean and covariance matrix of the training set examples’ latents and sample from a Normal distribution with those parameters.
- **Decoder:** As input, it receives the latent representation, and the AAs in the sequences sampled so far (or the whole sequence *x* during training). It comprises an embedding layer for the AA sequence, a transformer decoder stack (see Supplement **S.1.2**) and a head for AA prediction. We embed each sequence element with a learnt token embedding, a learnt positional embedding and a learnt next position embedding. The next-position embedding allows the prediction of protein sequences in arbitrary order (not only in N to C direction). Then, the decoder stack consists of multiple cross-attention blocks. These blocks can attend to the latent representation *z* and already generated tokens but, importantly for training, they cannot attend to not yet generated ones. The sampling is initialized with a start/stop token in the AA sequence. For training this is followed by the ground truth tokens, while during inference the predicted tokens are used (dashed line **Figure 1a**).

#### 2.3.2 VAE-training stage

The VAE‘s encoder is trained to map a protein sequence to a low dimensional latent representation, based on which the VAE‘s auto-regressive (AR) decoder aims to reconstruct the protein sequence. Training a VAE in conjunction with an AR decoder is challenging, due to “KL-vanishing”. To address this, we adopted the approach proposed by Li et al.[20] (see Supplement **S.1.1**). So, our training process (see **Figure 1b**) follows the steps below:

##### 1. Auto encoder (AE) training

We trained `CAPE-XVAE` like an AE for 500 epochs (KL divergence weight *β* in VAE loss function is zero). This focuses training solely on reconstructing the input sequence resulting in meaningful latent representations.

##### 2. Reset the decoder

Afterwards, the decoder is reset to its initialization. This counters the risk of the decoder learning faster than the encoder.

##### 3. Transition to VAE training

Over the next 250 epochs the KL-divergence weight in the loss function is increased linearly until it reaches one.

##### 4. VAE training

Then we continue training the model for an additional 750 epochs. This phase optimizes the full VAE objective with free bits. We fixed the free bits parameter *λ* as one.

To further increase the decoder’s reliance on the latent representation, we train the decoder to sample the protein sequence in random permutations of the ordering - not just N-C direction. This means that, while the normal next token prediction task is: here are amino acids 1, 2, 3 and 4 - please give me amino acid 5, we train the model on the following task: here are amino acids 22, 11, 59 and 37 - please give me amino acid 17. Further details about the hyper-parameter search, the eventually obtained hyper-parameters and the training procedure can be found in the Supplement **S.1.4**.

#### 2.3.3 Immune-visibility profile modification procedure

We modify the immune-visibility profile of sampled proteins by updating their latent representation using RL techniques. The RL problem is formulated as follows: `CAPE-XVAE`’s decoder represents the agent. The state *s*_*t*_ is the protein sequence produced so far. The decoder’s output distribution over all AAs for the next token represents the policy *π*. We also define an immune-visibility dependent reward function (see ‘Episode’ below). The episode ends when the decoder generates a stop token. Overall, the modification process follows the following steps (**Figure 1c**):

##### 1. Initialization

First a random *natural sequence x* (see `Box1`) is sampled from the dataset and a conditional latent distribution *p*(*·*|*x*) = 𝒩 (*µ*(*x*), *σ*(*x*)^2^) obtained from the encoder. From this an initial latent representation **z**^(0)^ is sampled. We restrict the modified latent representations to remain in a box around **z**^(0)^ by using an auxiliary latent representation **w** which is initially **0**. Changes to **w** then indirectly updates **z** according to Formula 1. This holds for each dimension *d* of **z** and each episode *j* (see below).

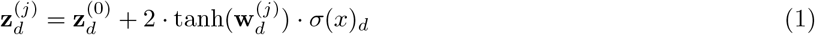

##### 2. Episode

At the beginning of each episode *j* the current auxiliary **w**^(*j*)^ is converted to a latent representation **z**^(*j*)^, which remains constant throughout the episode. The decoder uses this to sample a batch of 64 AA sequences in parallel in N-C direction. For each time-step *t* the decoder outputs a distribution over potential next AAs, representing the policy *π*. The temperature used to sample the next amino-acid *a*_*t*_ from this is geometrically decreased from 1.2 to 0.1 over the episodes of the modification phase. The gradient of the log-probability of the actually sampled AA with respect to the auxiliary (∇_*w*_ ln *π*_*θ*_(*a*_*t*_|*s*_*t*_)) is calculated and saved. Also, for each added AA the **reward function** is called and its result saved as this step’s reward *r*_*t*_. In addition to the current state (e.g. MGGKWSKSSSEAKWPAVRERMRRTEPADG), and the current action (e.g. V) the reward function receives the selected MHC-I alleles as input. The reward function penalizes or rewards presented peptides as soon as they are fully sampled. Most MHC-I epitopes have length 8, 9 or 10 AAs. Therefore, every time a new AA is sampled, the reward function calculates the *peptide reward* (see Subsection 2.5) for the terminal 8-, 9- and 10-mers of the combined sequence (e.g. MRRTEPADGV, RRTEPADGV, RTEPADGV). The reward function’s value *r*_*t*_ at the corresponding step in the trajectory is then the sum of these peptide rewards.

##### 3. Update latent representation

At the end of each episode *j*, the auxiliary latent representation **w**^(*j*)^ is updated to **w**^(*j*+1)^ using a version of the REINFORCE algorithm [21] (see Supplement **S.1.3**).

For each sampled position starting with the first and each of the 64 sequences generated in parallel, we update **w** according to the following formula 2:

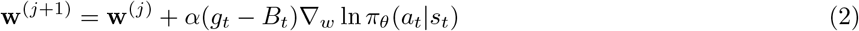

With:

α step size (we fixed this to 0.001)

*g*_*t*_ sum of future rewards 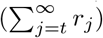

*B*_*t*_ average *g*_*t*_ across the batch

*s*_*t*_ the amino-acid sequence generated up to this position

*a*_*t*_ the amino-acid actually added to the sequence

ln *π*_*θ*_(*a*_*t*_|*s*_*t*_) the log-likelihood to sample *a*_*t*_ given state *s*_*t*_

∇_w_ the gradient with respect to **w**

Formula 2 demonstrates that when a particular action *a*_*t*_ is followed by rewards exceeding the average, it prompts an adjustment that boosts the probability of selecting that action. Conversely, if an action yields rewards below the average, the adjustment decreases the likelihood of selecting that action.

Then the algorithm goes back to step 2 until 100 updating steps have been completed.

### 2.4 CAPE-Packer

The underlying physics-based idea is, that proteins will fold into their energy minimizing state. So, `CAPE-Packer` searches for the energy minimizing sequence for a fixed 3D structure. By considering immune-visibility in the energy function (score function), the search process can be steered towards a desired immune-visibility profile. The implementation of `CAPE-Packer` builds upon the framework developed by Yachnin et al. [6] utilizing the *Rosetta Packer* [22, 23] (see 2.4.1) for protein design. Yachnin et al. modify Rosetta’s score function to discourage *Rosetta Packer* from incorporating MHC-II epitopes.

#### 2.4.1 Rosetta software for macro-molecular modeling and Rosetta Packer

Rosetta is a versatile software suite for modeling and analyzing protein structures. Next to others, it comprises algorithms dealing with structure prediction, docking and membrane proteins^3^. It has been developed for more than two decades and is extensively used in laboratories worldwide [24]. At its core lies the score (energy) function. This is the weighted sum of terms describing van der Waals energies, hydrogen bonds, electrostatics, disulfide bonds, residue solvation, backbone torsion angles, sidechain rotamer energies, and an average unfolded state reference energy [24].

Rosetta also features a key algorithm called the *Packer*, serving as a side-chain optimization tool. It identifies score function minimizing rotamers for a given sequence and fixed backbone structure (3D structure) [25]. Additionally, the *Packer* can be used for sequence design by considering different side-chain AA identities. In fact, this is its standard behavior and the *Packer* designs at every position, allowing for all 20 standard amino acid rotamers (position dependent subsets of eligible rotamers can be specified). The space of possible rotamer combinations expands rapidly, making enumeration infeasible. Therefore, Monte Carlo simulation is utilized. Starting from a random initialization, random changes are made to the rotamers. If the change decreases the score, it is always accepted. Conversely, if it increases the score, acceptance is stochastic with a probability that decreases with the energy [25].

#### 2.4.2 Rosetta Packer adaptations for immune-visibility modifications

Yachnin et al. [6] have suggested an additional term for the score function, that takes into account the number of peptides in the protein that get presented on the cell’s surface. In their case, they used MHC-II. We adapted their tool for MHC-I. The method only considers 9-mer peptides and out of the box it is only implemented using a PWM approach as well as a lookup database. While they mention that they have experimented with a tensorflow based presentation predictor, no implementation of this is known to us. In fact Mulligan [25] state that “In order to be compatible with the *Packer*, an energy term must either be residue-level pairwise-decomposable, or must otherwise be very fast to compute and update as rotamer substitutions are considered”. Therefore, we think the reason for the lack of a more advanced predictor backbone is, that the *Rosetta Packer* requires many score function evaluations - and, hence, peptide presentation evaluations. For a short 203 amino acid long Nef sequence, we counted roughly 42 million peptide evaluation requests. This must be multiplied by the number of considered major histocompatibility complex (MHC) alleles to arrive at the actual number of predictor calls. Using one of the modern and larger MHC-I presentation prediction state-of-the-art models - like *MHCflurry 2*.*0* [26] or *netMHCpan 4*.*1* [16] - is, therefore, very time intensive.

We implemented a new Rosetta class `MHCEpitopePredictorClient` that inherits from `MHCEpitopePredictor` and receives 9-mer evaluation requests from Yachnin et al.‘s framework. Our client then forwards these peptides together with the MHC-I alleles and the profile to modify towards to a server that we also implemented. This server returns the peptide’s reward (see Subsection 2.5). This number is multiplied by −1 and then integrated into the energy function used by the *Rosetta Packer* that ultimately designs the protein.

Being based on the *Rosetta Packer*, `CAPE-Packer` requires a 3D structure of the protein as input. *AlphaFold* [27] generated 3D structures of the *support* sequences (see `Box1`) were used for this. We allowed *Rosetta Packer* to sample from all AAs at every position in the sequence (we did not fix any regions of the protein).

### 2.5 Peptide reward

As the protein is designed, newly generated peptides by `CAPE-XVAE` receive a reward (see 2.3.3). Similarly, peptides sampled by `CAPE-Packer` receive a score modification (see 2.4.2) which is the negation of the reward. Their values depend on the targeted immune-visibility profile (see `Box2`). They are also influenced by how many of the considered MHC-I alleles present the peptide via the MHC-I pathway (*M*) and whether the peptide is present in the *natural sequences* or not (see `Box1`). The reward for a given peptide is computed as the product of the corresponding value in **Table 2** and *M*. The prediction of *M* is done using the PWM classifier for MHC-I presentation (see Subsection 2.1.2).

**Table 1:**
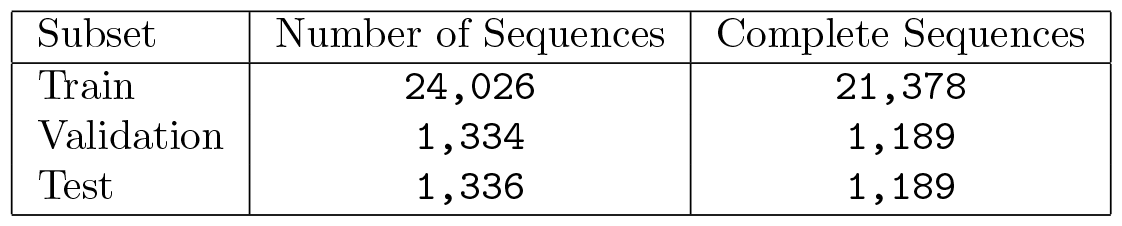
Number of unique sequences in dataset splits.

| Subset | Number of Sequences | Complete Sequences |
| --- | --- | --- |
| Train | 24,026 | 21,378 |
| Validation | 1,334 | 1,189 |
| Test | 1,336 | 1,189 |

**Table 2:**
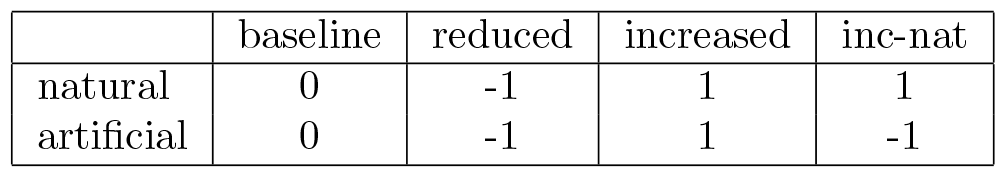
Rewards for a single peptide presentation: Rows signify if a peptide is present in the *natural sequences* or not. Columns represent the immune-visibility profile we aim to modify towards.

### 2.6 Evaluation

We assessed the generated sequences based on three criteria: immune-visibility, similarity to natural sequences and stability. Each criterion is detailed below. It is noteworthy that, in the absence of experimentally determined 3D structures, we employed *AlphaFold* [27] to predict the necessary 3D structures as needed.

#### 2.6.1 Visibility

To assess how MHC-I visible (see `Box2`) the proteins are to CTLs, we used *netMHCpan-4*.*1* [16]. We used the same MHC-I alleles during the evaluation step as we had used for the modification process. The visibility number is calculated as outlined in `Box2`.

#### 2.6.2 Similarity

We assess the similarity of the generated sequences to natural ones with regards to three criteria: local-, structural- and functional similarity.

- **Local similarity** is used to demonstrate the extent to which local sequence patterns in protein design are captured. To assess whether there is a similarity in parts of the sequences to the natural sequences (see `Box1`), we checked short sub-sequences of length k (k-mers) in the sequence against all k-mers occurring in the natural sequences. The percentage that is also present in the natural sequences we define as the *k-mer similarity*. 9-mer similarity is of particular interest since most presented peptides have 9 AA length, making it especially relevant for vaccine design, where we try to design sequences that more broadly capture epitopes across variants of a protein. To assess the overlap between k-mers we define the following metrics: (a) the *k-mer recall* of a query sequence (a generated or natural sequence) wrt a subject sequence (a natural sequence) is the percentage of k-mers in the subject sequence that are also present in the query sequence; (b) the *k-mer precision* of a query sequence wrt a subject sequence is the percentage of k-mers in the query sequence that are also present in the subject sequence; (c) for a specific k-mer, *k-mer precision* is defined as the percentage of the natural sequences that include this k-mer.
- **Structural similarity** is used to compare the overall fold of the predicted structure of the designed sequence to that of its *closest support sequence* (see `Box1`). The TMscore between the two structures was calculated using *TMalign* [28]. The TMscore ranges between zero and one. A score of one signifies a perfect match while scores below 0.2 are associated with randomly chosen unrelated proteins. Scores higher than 0.5 assume generally the same fold in SCOP/CATH ^4^.
- **Functional similarity**: Inspired by the idea in Generative Adversarial Networks (GANs), where examples are judged to be “better” if a discriminator has a harder time to distinguish between actual and generated examples, we train a classifier to distinguish between naturally occurring and generated sequences. We want to focus on functional aspects and leverage the capabilities of third party tools. So, the first step in our classification pipeline is the GO term predictor of Boadu et al. [29] that - based on a sequence’s *AlphaFold* generated pdb file - predicts scores for `11,156` Molecular Function GO terms. After normalization, we performed a PCA keeping the first 19 components which cover 99% of the observed variance. Eventually, a Gaussian Naive Bayes classifier (called *GO based Sequence Pack Classifier*) is trained to classify the sequences into 8 different origins: support, `CAPE-XVAE`-*baseline*/*reduced* /*increased* /*inc-nat*, `CAPE-Packer`-*baseline*/*reduced* /*increased*. As we do not want to perform prediction but simply want to assess the classifier’s ability to distinguish between the classes, no holdout dataset was used.

#### 2.6.3 Stability

To assess the stability of our generated protein sequences we select a subset of these and the support sequences and run a Molecular Dynamics (MD) simulation on this. We then calculate the maximum root mean squared distance (RMSD) (based only on CA-atoms) between the confirmations observed during the simulation and the original *AlphaFold* prediction. We use OpenMM to run the MD simulation for `2` microseconds (`2` femtoseconds step-size) at a temperature of `25` degrees Celsius using the amber99sb force field and an implicit solvent model (amber99 obc). Positions were reported every 4 picoseconds. Due to the high computational requirements, we only performed this analysis on a sub-sample of sequences. Within each immune-visibility profile, these are randomly selected.

### 2.7 2D sequence space visualizations

We want to produce a 2D figure illustrating the similarity between sequences. The idea is to start by representing each sequence by a vector in which each dimension represents the dissimilarity (see below) between the represented sequence and one of the other sequences. Owing to the infeasibility of comparing each sequence with all other sequences, we compare each sequence only to the 300 *support sequences* (see `Box1`). Afterwards t-SNE reduces this to a 2D representation.

We compute dissimilarity based on the pairwise alignment score of the sequence with the *support sequence*. Gotoh global alignment algorithm [30] as implemented by the biopython [31] PairwiseAligner (using BLOSUM62, open gap score of `-20` and extend gap score of `-10`) was used for this. The dissimilarity is then defined as the difference between the alignment score of the sequence with itself and the alignment score of the sequence with the *support sequence*. To further reduce the dimension of these vectors to only two, we use t-SNE with a perplexity of 30 and running for 500 iterations. We have observed similar results for a broad range of number of support sequences and perplexity numbers.

## 3 Results and discussion

In this section, we present the outcomes of the experiments conducted using `CAPE-XVAE` and `CAPE-Packer` for modifying immune-visibility concerning six common MHC-I alleles from a hypothetical patient: HLA-A*02:01, HLA-A*24:02, HLA-B*07:02, HLA-B*39:01, HLA-C*07:01, HLA-C*16:01. Evaluation criteria include MHC-I immune-visibility (see `Box2`), similarity to natural sequences, and the stability of the generated proteins. For each *visibility profile* (see `Box2`), 100 `CAPE-XVAE` and 300 `CAPE-Packer` sequences were generated.

### 3.1 Differential distribution in sequence space: CAPE-XVAE sequences distribute more naturally than CAPE-Packer ones

We first explored the landscape of naturally occurring and `CAPE-XVAE` and `CAPE-Packer` generated sequences (**Figure 2**, see Subsection 2.7). Notably, the baseline sequences (see `Box2`) produced by `CAPE-XVAE` are dispersed across the natural sequence space in **Figure 2a**, showing no aggregation in any particular region. However, the modification step removes them from several areas of the sequence space and moves some even outside the range occupied by natural sequences. This is not observed for `CAPE-XVAE`-*inc-nat* sequences.

**Figure 2:**
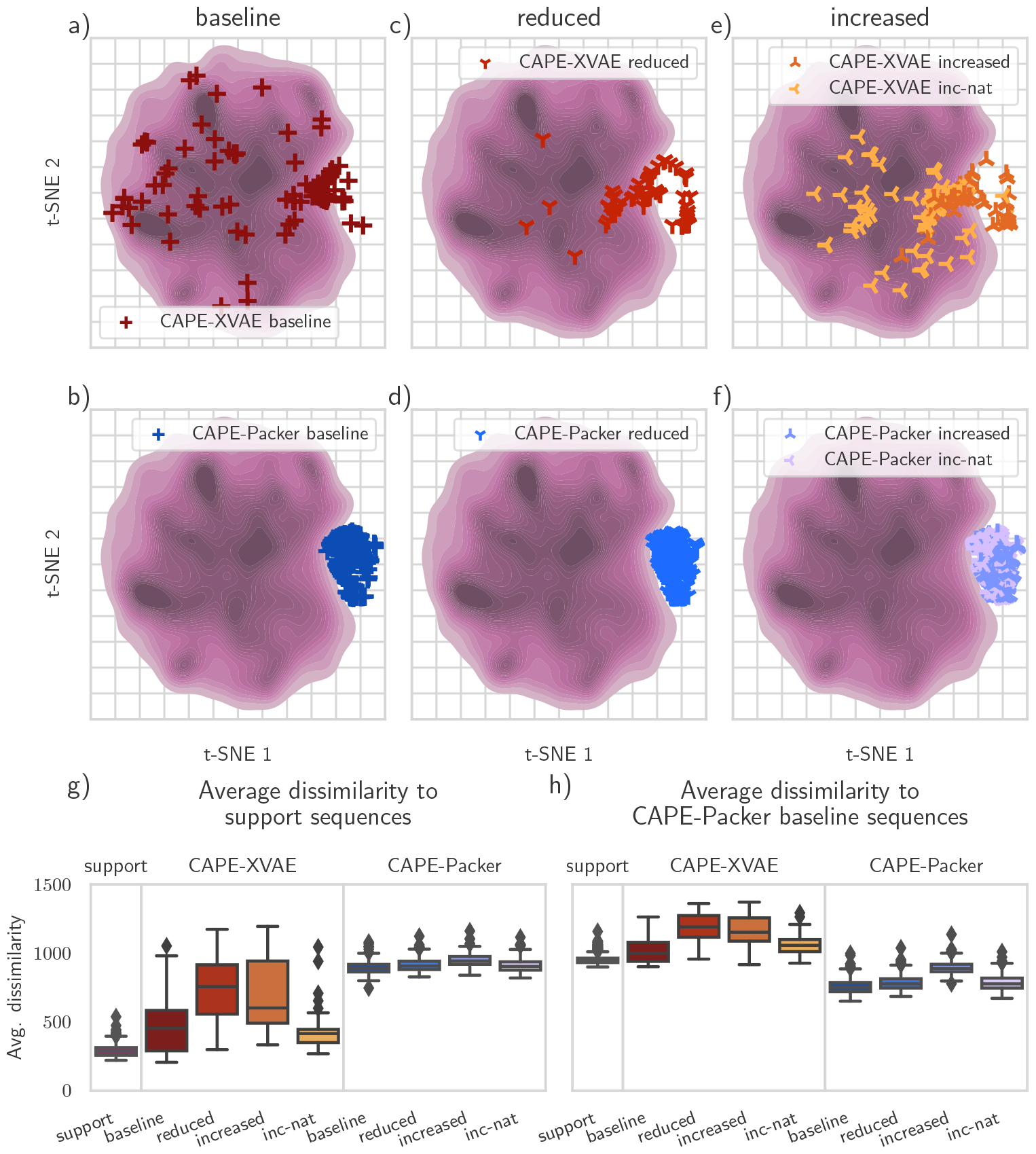
Differential distribution of CAPE-XVAE and CAPE-Packer sequences in natural sequence space. The density of the natural sequences used to train the models is depicted pink in the background. (a,b) The dark red/blue crosses correspond to the sampled sequences obtained through `CAPE-XVAE`/`CAPE-Packer` without modification process (baseline). (c,d) Show the reduced immune-visibility results. The red/blue crosses are the finally obtained `CAPE-XVAE`-*reduced* /`CAPE-Packer`-*reduced* sequences. (e,f) Here increased and increased-natural immune-visibility results are displayed. `CAPE-XVAE` designs are in orange and `CAPE-Packer` ones in blue. The light/dark crosses show the finally obtained *inc-nat* /*increased* sequences. (g,h) The distribution of the average dissimilarity to the 300 support sequences (g, see `Box1`) as well as to the `CAPE-Packer`-*baseline* sequences (h). Colors correspond to the other plots

Starkly, the sequences generated by `CAPE-Packer` all aggregate outside the natural background distribution of sequences (**Figure 2bdf**). Despite appearing close in t-SNE plots (**Figure 2a-f**), which is a cluster-preserving but not distance-preserving dimension reduction technique, the sequences outside the natural background distribution are highly dissimilar from each other. In fact, **Figure 2h** illustrates that the `CAPE-Packer` baseline sequences are markedly different from all other sets and even differ from each other, while support sequences are similar to themselves (**Figure 2g**).

### 3.2 CAPE-XVAE designs incorporate many natural peptides

In **Figure 3** we compare the generated sequences to naturally occurring ones with respect to immune-visibility (**Figure 3a**, see `Box2`), local similarity (**Figure 3b-e**) and structural similarity (**Figure 3f**, see 2.6.2). These results indicate that `CAPE-XVAE` and `CAPE-Packer` are effective in altering MHC-I immune-visibility (**Figure 3a, Table 3**) while maintaining structural similarity.

**Table 3:** Mean Visibility: Rows represent the sequence sources and columns the modification profile (see `Box2`)

| mean visibility | baseline | reduced | increased | inc-nat |
| --- | --- | --- | --- | --- |
| Support | 87 |  |  |  |
| CAPE-XVAE | 86 | 52 | 146 | 96 |
| CAPE-Packer | 71 | 41 | 286 | 42 |

**Figure 3:**
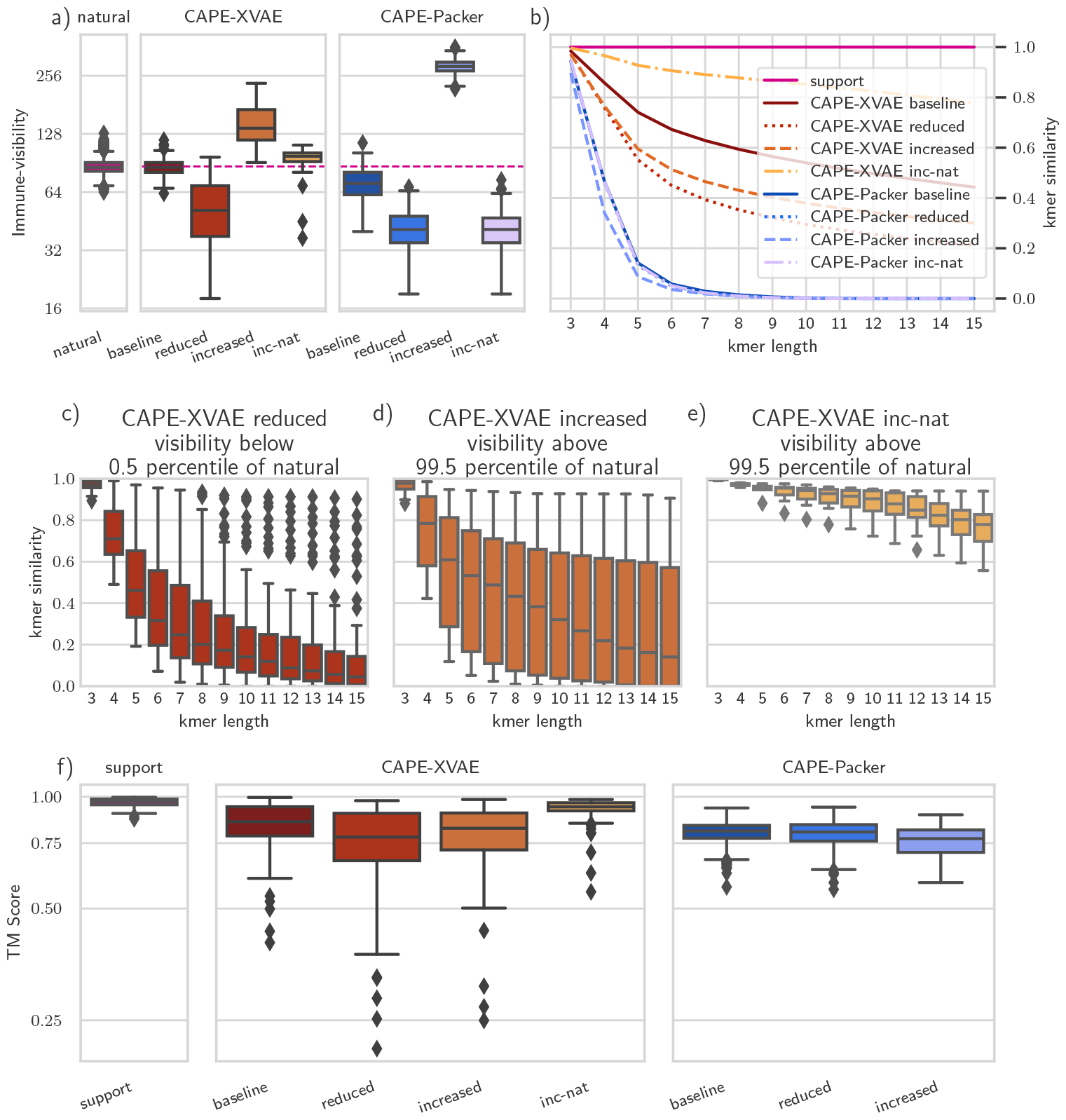
CAPE-XVAE alters MHC Class I immune-visibility while maintaining natural sub-sequence characteristics. (a) **Visibility:** Shows the distribution of the visibility of the various sequence sets. The visibility is calculated equivalently to how the *reward function* is calculated for the *increased* modification profile (See Subsection 2.3.3). The dashed pink line signifies the mean of the natural sequences. (b) **K-mer Similarity:** Dependent on k-mer length, this chart shows the average proportion of k-mers with that length in the sequences that can also be found in the natural sequence data. One means that all k-mers can be found in the data and zero means that none of the k-mers can be found in the data. (c-e) Boxplots show the k-mer length dependent distribution of k-mer similarity for the `CAPE-XVAE` modified sequences that are either in the bottom (*reduced*) (c) or top (d,e) 0.5 percentile (*increased, inc-nat*) of natural sequences. (f) Shows the distribution of TM Score as calculated by *TMalign* [28] between the sequences and their *closest natural sequence*. Color scheme continues from **Figure 2**

In **Figure 3a** we see that `CAPE-Packer` can produce extremely immune-visibility modified sequences. `CAPE-Packer` is only cognizant of protein fold, without access to the natural grammar of protein function. It seems unlikely, that longer natural peptides are randomly sampled during *Rosetta Packer’s* Monte Carlo simulation process (see Subsection 2.4), preventing it from incorporating those (**Figure 3b**) even when rewarded - like in the case of `CAPE-Packer`-*inc-nat*. This limits `CAPE-Packer`’s usefulness in vaccine design. In fact, `CAPE-Packer`-*inc-nat* is effectively turned into `CAPE-Packer`-*reduced* (we think this is because it is very unlikely to sample actually occurring visible natural 9-mers, which are the only ones treated differently between the two visibility-profiles) and will, therefore, be ignored from here on. With regards to `CAPE-Packer`-*reduced*, Yachnin et al. [6, Figure 2b] observed a slightly higher reduction in MHC-II visibility for deimmunizing human erythropoietin when using the same weighting of the immune-visibility term in the score function as us (see 2.4). With regards to structural similarity, **Figure 3f** shows that, despite their unnatural looking sequences, there are `CAPE-Packer` proteins with high TMscores.

In comparison, `CAPE-XVAE` sequences tend to be less immune-visibility modified (**Figure 3a**). The incorporation of artificial visible peptides enables `CAPE-XVAE`-*increased* to achieve higher visibility than `CAPE-XVAE`-*inc-nat*. **Figure 3b** demonstrates that nudging `CAPE-XVAE`-*inc-nat* to only incorporate natural visible peptides also seems to cause it to incorporate more of the natural k-mers in general.

The analysis of k-mer similarity for *highly* and *lowly immune-visible* (see `Box2`) `CAPE-XVAE` generated sequences reveals that, even within strongly modified sequences, many exhibit high k-mer similarity to natural sequences (**Figure 3c-e**). In particular `CAPE-XVAE`-*inc-nat* designed sequences demonstrate high values. This conservative behavior of `CAPE-XVAE`-*inc-nat* may be beneficial in vaccine design, which might benefit from representative sequences that sample the breadth of potential MHC-I epitopes (visible 8, 9 and 10-mers) observed in the pathogen. Even for generated sequences with reduced visibility, remaining close to natural proteins in sequence space may increase the chances that complex nuances of protein function are retained.

The structural similarity of `CAPE-XVAE` designs is more dispersed than for `CAPE-Packer` designs (**Figure 3f**). Very interestingly, `CAPE-XVAE`-*inc-nat* designs tend to boast higher TMscores than `CAPE-XVAE`-*baseline* ones. Applying some form of RL inspired latent space search might, therefore, be beneficial in the protein design process of VAE approaches in general.

### 3.3 There are tradeoffs between modifying immune-visibility and maintaining local and structural similarity

Modifying immune-visibility comes at a cost: the more the sequences diverge, the more they lose local similarities to the training set, though their stability remains unaffected. **Figure 4** demonstrates, that the further we move away from the data distribution, the less similar the generated sequences become at the peptide length pertinent to antigen presentation (9-mers). Stability does not seem to be affected as assessed using a sub-sample of RMSDs from computationally expensive MD simulations. Despite this trade-off, we find among `CAPE-XVAE` designs many that contain significant 9-mer similarity even with high visibility. In contrast, 9-mer similarity is not present for any of the `CAPE-Packer` generated sequences. For `CAPE-XVAE`, the proportion of high TMscore (above 0.8) proteins generated decreases with distance to the natural visibility range, while this seems not to be the case with `CAPE-Packer`.

**Figure 4:**
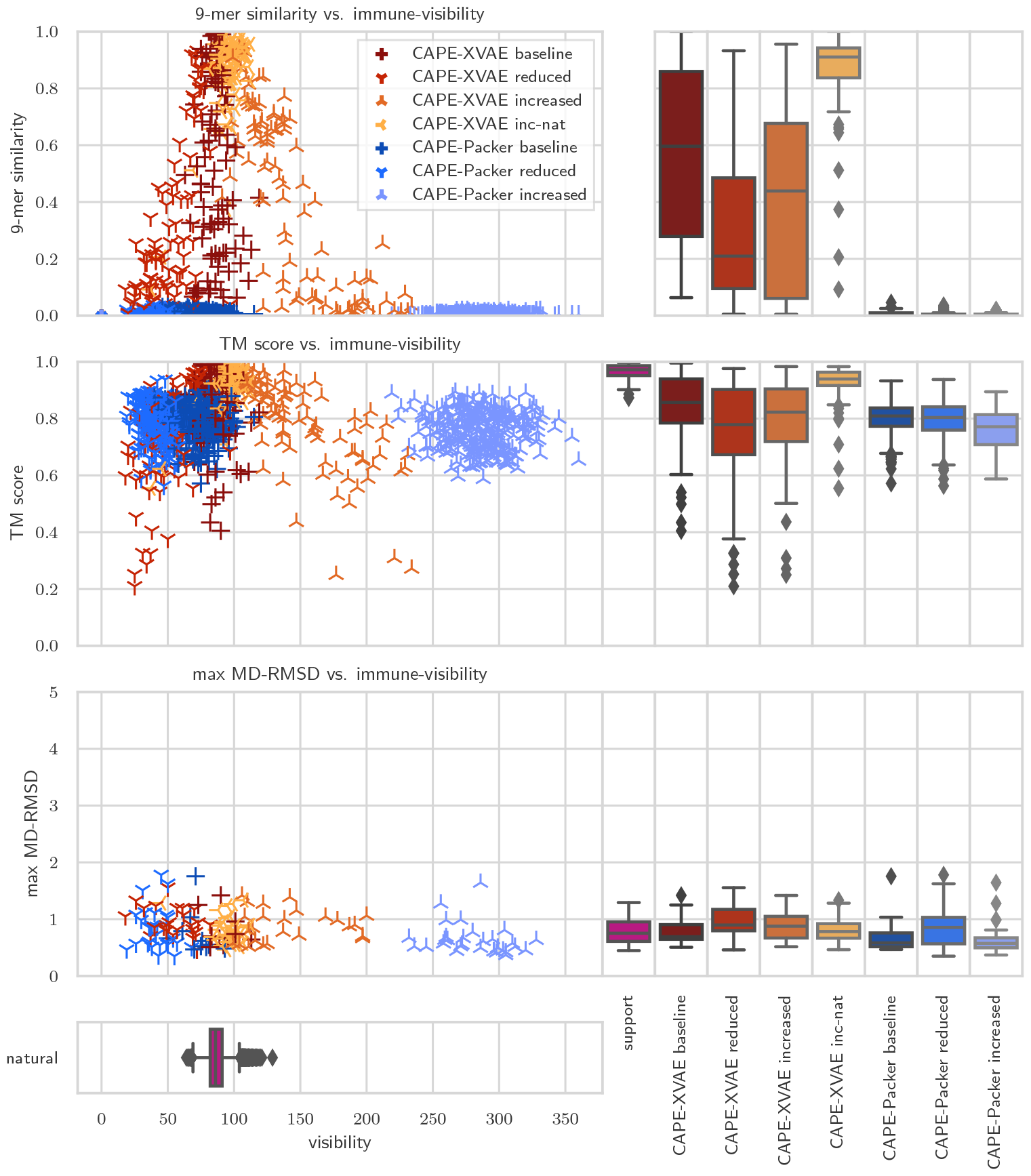
Assessing similarity and stability dynamics in protein design: A comparative study of visibility-dependent trends in CAPE-XVAE and CAPE-Packer generated sequences. In each row of this chart, we show how the natural data sequences compare to `CAPE-XVAE` and `CAPE-Packer` generated sequences with respect to one of the following criteria: 9-mer similarity, TM Score and maximum RMSD in an MD simulation. Each dot in the left column represents a single sequence - the same color coding as in **Figure 2** applies. The x-axis shows the immune-visibility. The three boxplots per row on the right display the distribution within the compared values for the support as well as the two model generated sequences. The horizontal boxplot on the bottom shows the distribution of visibility within the data.

### 3.4 Distinguishing between designed and natural sequences is difficult based on predicted function

We explored whether `CAPE-XVAE` and `CAPE-Packer` designs are likely to retain natural functionality (see *Functional similarity* 2.6.2). For each sequence we first predicted the scores for various molecular functions. This long vector representation for each sequence was reduced by principal components analysis. The first two components **Figure 5** (explaining `68%` of variance) visualize the challenge posed to distinguish many of the generated sequences from the support sequences. Certainly, many `CAPE-XVAE` generated sequences would be challenging to distinguish this way from natural sequences.

**Figure 5:**
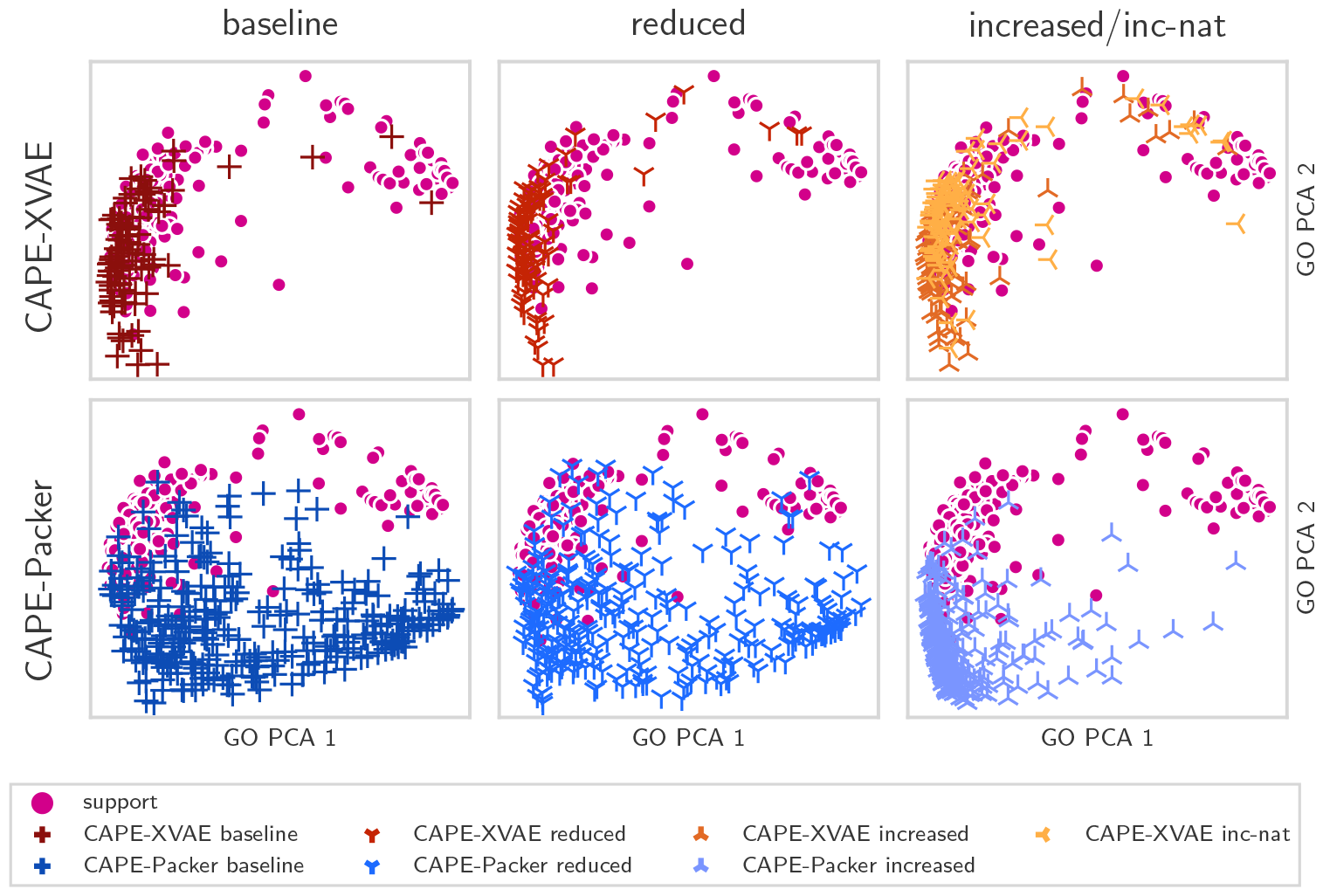
Many designed sequences are hard to distinguish from support sequences based on molecular function GO term predictors. Each of the six plots shows the first two principal components used to train the *GO based Sequence Pack Classifier* (see Subsection 2.6.2). The figure compares in two lines the results for `CAPE-XVAE` and `CAPE-Packer` designed sequences (as colored crosses) to always the same support sequences (pink). Each column depicts a different visibility modification profile.

We then trained a *GO based Sequence Pack Classifier* (Subsection 2.6.2) using the first 19 principal components covering 99% of variance. This classifier also struggles to distinguish between the various sources as demonstrated in the confusion matrix (**Table 4**). `CAPE-Packer` sequences are easier to distinguish from natural sequences than `CAPE-XVAE` generated sequences where only 60% of support sequences are correctly identified as support sequences, while it considers 28% of them to be `CAPE-XVAE`-*baseline* sequences. It also classifies 28% of `CAPE-XVAE`-*baseline* as well as many `CAPE-XVAE` modified sequences as support sequences. In fact, a third of the `CAPE-XVAE`-*inc-nat* sequences are classified as natural.

**Table 4:**
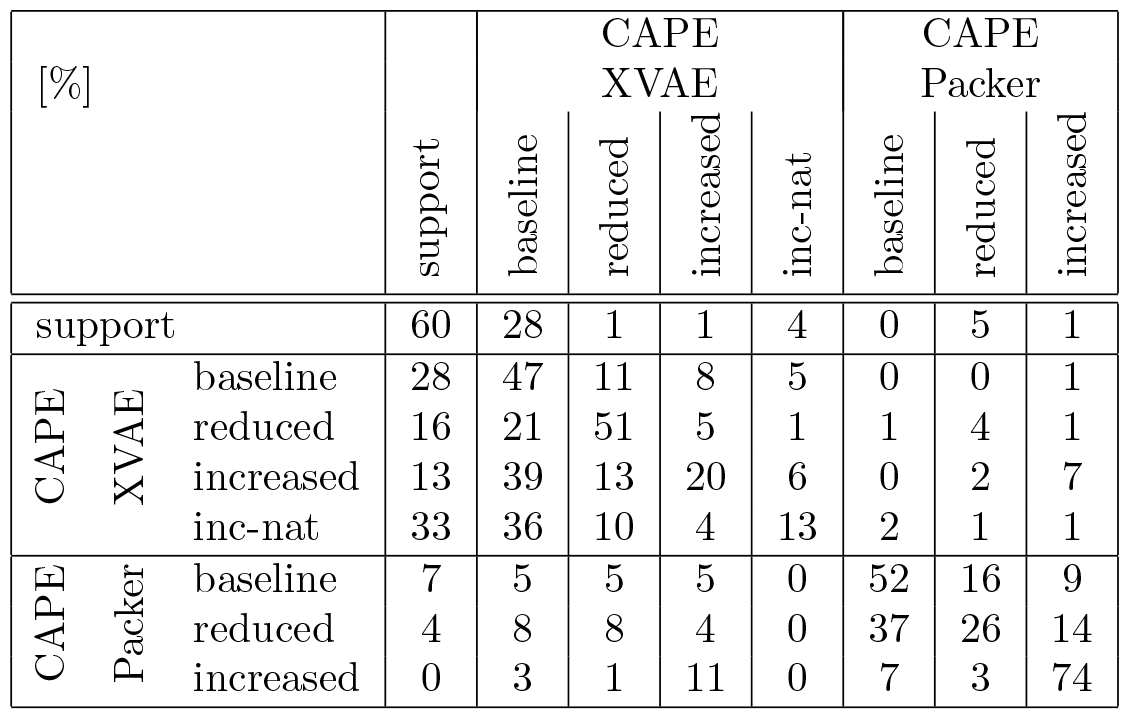
Confusion matrix: The rows signify the actual sequence origin, while the columns their predicted one

**Table 5:** Hyper-parameter search

| Hyper-parameter | Distribution | Selected |
| --- | --- | --- |
| Latent variables in a latent representation ($L$) | 1 | 1 |
| Dimension of a latent vector ($l$) | 32 | 32 |
| Lambda ($\lambda$) | 1.0 | 1.0 |
| Model dimensions ($d$) | 256 | 256 |
| Feed-forward dimensions | 512 | 512 |
| Number of heads | 8 | 8 |
| Number of layers | 1 to 5 | 3 |
| Dropout | Uniform(0, 0.4) | 0.015 |
| Learning rate | Log-Uniform(0.0001, 0.01) | 0.00013 |
| Batch size | 32 | 32 |

It appears that as a whole there are differences in predicted function between natural and generated sequences. More importantly however, a considerable proportion of the proposed `CAPE-XVAE` sequences closely resemble natural ones in terms of predicted molecular function. In practical applications, these sequences may be more desirable to advance.

### 3.5 CAPE-XVAE-*inc-nat* designs tend to have high mean recall and precision

While the intricacies of vaccine design are best managed by vaccinologists with the capacity to experimentally test various strategies, `CAPE-XVAE` offers a tool to suggest chimeric sequences capable of exposing the immune-system to a representative set of epitopes, potentially enhancing population-wide protection. As `CAPE-XVAE` proposes similar sequences to natural ones, we were interested in understanding how diversely potential epitopes (k-mers predicted to be immune-visible) were sampled from the natural sequences. **Figure 6** shows how the natural sequences compare to `CAPE-XVAE`-*inc-nat* designed sequences with respect to how well they recall naturally occurring visible peptides. First, we observe that even the sequences with the highest recalls in expectation include a third of the visible peptides of a randomly chosen natural sequence. Furthermore, we find that although the highest recall value can still be found in naturally occurring sequences, the set of designed ones has a significantly higher average mean recall as determined by a two-sided t-test. Interestingly, for highly visible sequences, this is even more pronounced. In conclusion, we believe proteins designed using language model (LM) can be valuable constituents of vaccines due to their ability to tap into the array of natural viral variant epitopes - even for less common ones.

**Figure 6:**
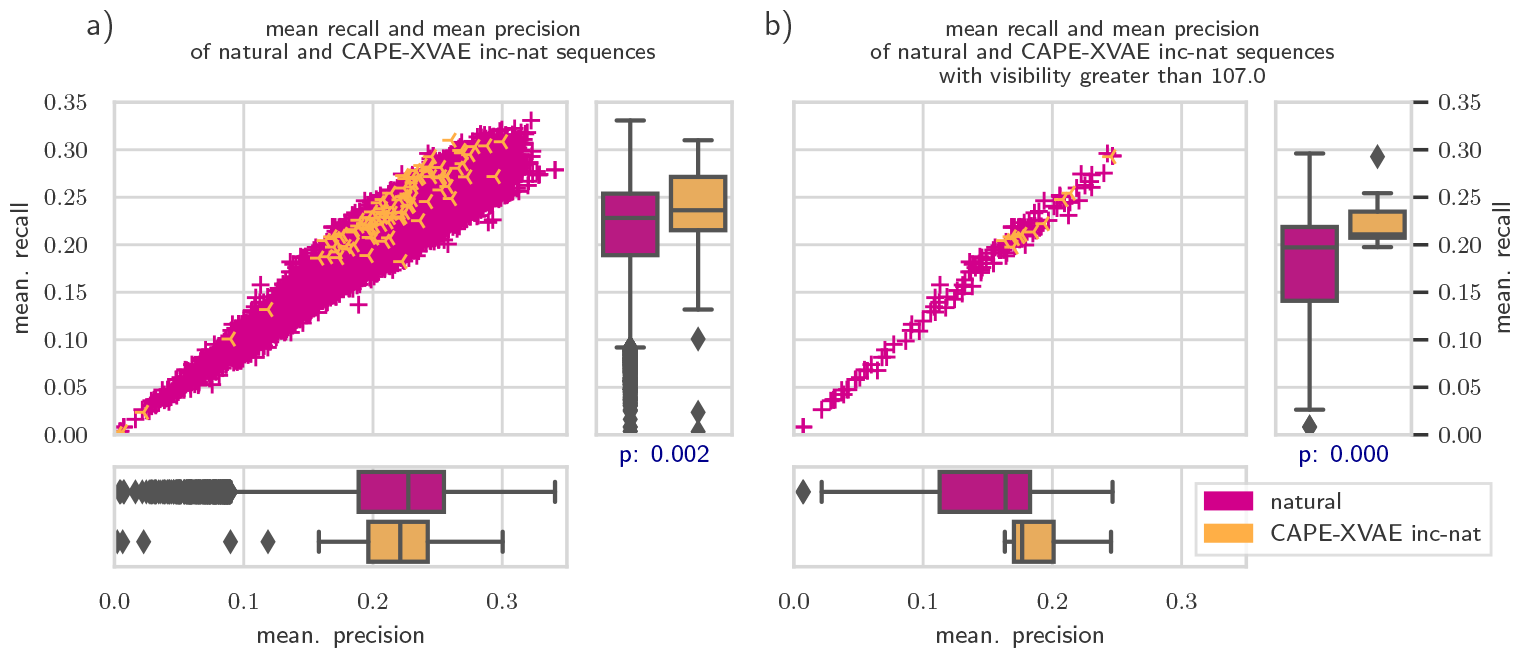
Average recall and precision of natural and CAPE-XVAE-*inc-nat* designed sequences. Each dot in the scatterplots represents a natural or designed sequence. Its x-coordinate shows the sequences average precision over all natural sequences, while its y-coordinate depicts the sequences average recall over all natural sequences (8-, 9- and 10-mers). The boxplots on the right and bottom depict the distributions of average recall and precision for the two sources. The blue p-values are the t-test p values for the null hypothesis that the mean recalls of natural and designed sequences are the same. While plot (a) depicts the situation considering all sequences, (b) is restricted to highly immune-visible sequences (see `Box2`)

**Figure 7** then explores the ability of the highest mean recall `CAPE-XVAE`-*inc-nat* sequence with a TMscore of above 0.95 to capture the naturally occurring k-mer and epitope diversity. We find that all visible 9-mers in the `CAPE-XVAE` designed sequence can indeed be found in the natural sequences (**Figure 7b**). Only 3 of the visible 9-mers (red boxes in **Figure 7a**) were present in less than 0.1% of the natural sequences (**Figure 7b**). In contrast, six of these were present in more than 50% of all natural sequences. Finally, **Figure 7c** shows the distribution of the visible 8, 9 and 10-mer recall. We find that only a few sequences have less than 10% of their visible peptides being present in the designed sequence.

**Figure 7:**
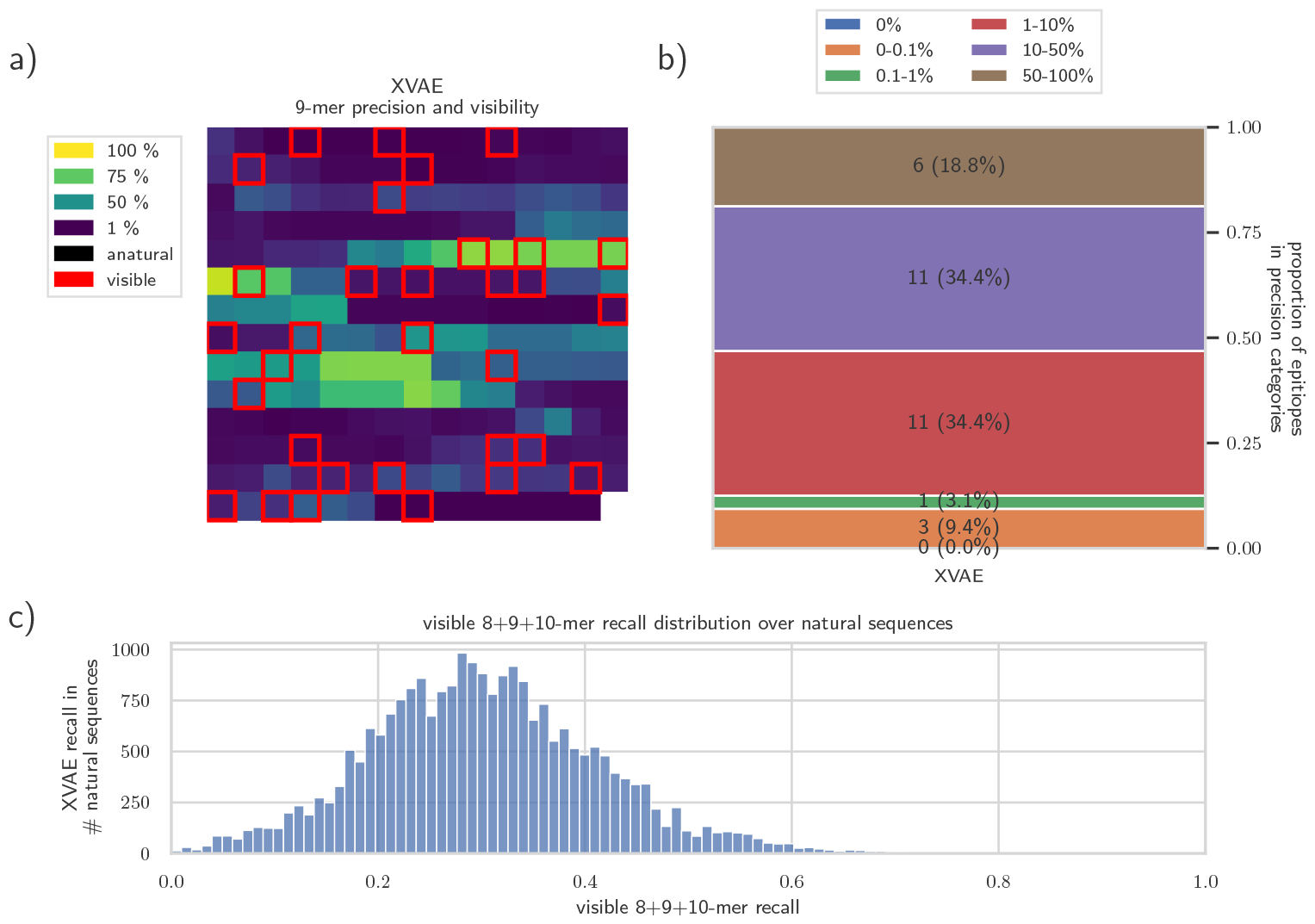
Language models in vaccine design: borrowing from the diversity of existing protein variants. `CAPE-XVAE` supports the design of vaccines that contain real epitopes existing across variants of a virus. This figure shows an epitope map of the highest mean recall `CAPE-XVAE`-*inc-nat* designed sequence (see **Figure 6**). Its predicted visibility is 103 and its mean recall 31%. The *GO based Sequence Pack Classifier* predicts it to be a *support* sequence. We investigate where each epitope is derived from within the natural distribution of input sequences. (a) We present 9-mer peptides in the designed Nef protein sequence using a sliding window approach advancing 1 amino acid at a time. The N-Terminal 9-mer is top left progressing one amino acid at a time to the C-Terminal 9-mer on the bottom right. The colors signify how often these 9-mers occur among the natural sequences (see *k-mer precision* in 2.6.2). A red box signifies that this 9-mer is predicted to be presented by at least one MHC allele (a visible 9-mer). (b) The labels of the stacked bar chart are the number of visible 9-mers [red boxes in (a)] in a precision category (signifying the percentage of natural sequences that contain this visible 9-mer). (c) For all natural sequences we identify the visible 8-, 9- and 10-mers and calculate the percentage of those present in the designed sequence. The histogram shows the distribution of this number over all natural sequences (see *k-mer recall* in 2.6.2).

### 3.6 Limitations

The results presented are limited by several factors. First, we are heavily reliant on the state-of-the-art *AlphaFold* folding prediction model to produce accurate structures. The assessment of structural and functional similarity of the designed sequences to naturally occurring ones relies on these. *AlphaFold* was trained on naturally occurring sequences - in particular for `CAPE-Packer` designs, which are very divergent sequences, it is questionable whether predicted structures will remain accurate - and if they even were able to fold into those.

Next, we are using a GO term predictor to assess functional similarity. On occasions, even a single mutated amino acid can lead to a loss of function. GO term predictors that are trained to predict the function of working proteins may miss those subtle differences. However, we are not aware of any ML method in protein design that performs a major modification to a protein and still proposes a functional protein in each attempt. In reality, the protein design process involves many filtering steps. The *GO based Sequence Pack Classifier* can be seen as another way to filter sequences. The difficulty of assessing designs is a challenge for the whole field and ultimately only confirmation in the wet lab on the output of tools like ours will deliver definite results.

Finally, our model does not directly control immune-reaction by CTLs, but only visibility to CTLs (see 2.1.1). Furthermore, other immune-system components (e.g. Abs) were not considered.

## 4 Conclusions

Designed protein-based medicines hold transformative potential for healthcare, serving as potent vaccines and therapeutics for various diseases. Ensuring these medicines do not inadvertently trigger an immune-response, except when intended (as with vaccines), is crucial. This paper focuses on the design of proteins with controlled visibility to CTLs via the MHC Class I pathway.

We introduce `CAPE-XVAE`, a latent space controlled language model trained on natural sequences to propose new proteins that are more visible to the immune-system when needed, and stealthy when not. We compare its outputs to those generated by another method we developed - the physics-based `CAPE-Packer` which is based on the deimmunization protocol developed by Yachnin et al. [6] and ignorant of the target protein’s natural sequence space.

We find that both methods can modify immune-visibility compared to natural sequences while maintaining high levels of structural similarity. As `CAPE-Packer` is not limited by the diversity in the target protein’s natural sequence space, we find that it is more flexible in adjusting immune-visibility. However, designing proteins within the boundaries of natural sequence space is crucial for two main reasons. Firstly, it helps in preserving the known and potentially undiscovered functions of the protein being designed by conserving local functional constraints. Secondly, for vaccines, it allows to incorporate the diversity of in particular sequence-based MHC-I epitopes. That `CAPE-XVAE` can capture this local sequence similarity is demonstrated by the substantial proportion of naturally occurring short sub-sequences within its generated sequences (k-mer similarity). In fact, it generated sequences with particularly high visible peptide recall, allowing it to capture the diversity of potential epitopes found in viral protein sequence variants. This contrasts with `CAPE-Packer`, which is unable to incorporate those.

We conclude that methods like `CAPE-Packer` that disregard sequence information will be useful when designing novel proteins under immune-visibility constraints. The designer needs to have full knowledge of the functions the protein is supposed to fulfill as well as the mechanism to achieve those. Functional regions can then be fixed, before allowing the algorithm to design around them. However, when we want to modify naturally occurring proteins where no full information about functionality and its mechanism is available, the hope is for machine learning algorithms to infer these from countless existing examples.

Flexible models trained from natural sequences and able to incorporate naturally occurring epitopes, like `CAPE-XVAE`, will be particularly useful for vaccine design. `CAPE-XVAE` can be easily adapted to modify for a broad range of immune-visibility profiles (see `Box2`) dependent on the latest developments in vaccine design. Generating chimeric sequences could potentially make escape mutations more unlikely and capturing the diversity in naturally occurring sequences may provide better, more universal vaccines in the future, compared to those that use just one strain of a virus. The concept will need validation, and this will be the subject of future studies.

However, with regards to reducing immune-visibility, we have also observed that `CAPE-XVAE` is not able to completely hide proteins from CTLs in our example case. As it solely relies on the distribution of natural protein sequences it might struggle to propose sub-sequences that need to be introduced to achieve invisibility if they do not exist in the training data. Because of this, we think that new ML-based approaches will need to be developed implementing a tunable and hybrid approach. These methods would incorporate structural information in addition to sequence. They could also utilize information from other proteins in addition to the target protein and be able to diverge by a defined and tunable amount in sequence space. Eventually, more complex immune-visibility profiles - considering immunogenicity and immunodominance also with regards to other components of the immune-system (e.g. MHC-II) - need to be explored, but using a feedback loop with wet-lab validation.

## Acronyms

AA: amino acid
Ab: antibody
AE: auto encoder
AR: auto-regressive
CAPE: Controlled Amplitude of Present Epitopes
CTL: Cytotoxic T-lymphocyte
GAN: Generative Adversarial Network
LM: language model
MD: Molecular Dynamics
MHC: major histocompatibility complex
MHC-II: MHC Class II
MHC-I: MHC Class I
ML: machine learning
PWM: position weight matrix
RL: reinforcement learning
RMSD: root mean squared distance
TCR: T-cell receptor
VAE: Variational Autoencoder

## Acknowledgements

This work was supported by the United Kingdom Research and Innovation (grant EP/S02431X/1), UKRI Centre for Doctoral Training in Biomedical AI at the University of Edinburgh, School of Informatics. For the purpose of open access, the author has applied a creative commons attribution (CC BY) licence to any author accepted manuscript version arising. This project was supported by the Royal Academy of Engineering and the Office of the Chief Science Adviser for

National Security under the UK Intelligence Community Postdoctoral Research Fellowship programme.

J.A. Alfaro was supported by (i) United Kingdom Research and Innovation (grant EP/S02431X/1), (ii) the project ‘International Centre for Cancer Vaccine Science’ which is carried out within the International Agendas Program of the Foundation for Polish Science, cofinanced by the European Union under the European Regional Development Fund. The authors would like to thank ‘CI-TASK, Gdansk’, and the ‘PL-Grid Infrastructure, Poland’ for providing their hardware and software resources. Dr. Rajan was supported by the KATY Consortium H2020-SCI-FA-DTS-2020-1.

## 6 Author contributions

**Hans-Christof Gasser:** Conceptualization, Methodology, Software, Validation, Writing - Original Draft, Writing - Review & Editing, Visualization **Diego Oyarzun:** Conceptualization, Writing - Review & Editing, Visualization, Supervision **Ajitha Rajan:** Conceptualization, Writing - Review & Editing, Visualization, Supervision **Javier Alfaro:** Conceptualization, Writing - Review & Editing, Visualization, Supervision

During the preparation of this work the author(s) used ChatGPT in order to increase the readability of the text. After using this tool/service, the author(s) reviewed and edited the content as needed and take(s) full responsibility for the content of the publication.

## S Supplementary material

### S.1 CAPE-XVAE

#### S.1.1 Variational Autoencoders

The basic idea behind AEs and VAEs is, that there are some hidden “latent” variables that can be used to reconstruct high dimensional data from a lower dimensional latent representation. For pictures of faces, these latent variables could be the length, color and shape of hair, the facial expression, etc.

AEs consist of an encoder that maps an input example *x* to a latent representation vector *z* and a decoder that takes in this vector and reconstructs the example. They are trained together to minimize the reconstruction error. Unfortunately, AE may suffer from “holes” in the latent space where there are no corresponding realistic examples. VAEs aim to address this by incorporating an additional KL-divergence term into their objective function, that encourages the latent representations to be normally distributed around the origin in latent space.

Unfortunately, VAEs may suffer from another problem known as “posterior-collapse” or “KL-vanishing”. This arises when using a very powerful decoder - often an AR model. In those cases, the following behaviour often ends up being optimal to learn for the model. The encoder simply learns to minimize the KL-divergence term while the decoder ignores the latent representations and directly learns to construct realistic examples. Various methods have been developed to mitigate it and preserve the meaningfulness of the latent representations. We follow an approach suggested by [20] which combines three methods.

- **Free bits:** The KL divergence term in the objective function is cut off by a hyper-parameter *λ* in each latent dimension. This allows for a controlled amount of information about the input to be retained in the latent representation without penalty.
- **Annealing:** During training, the KL-divergence term in the objective function gets weighted by a factor *β*. This starts out at zero - effectively allowing AE training for a certain period of time - before slowly rising to one.
- **Resetting the Decoder:** The uninformative latent representations initially provided by the encoder during training may lead to the decoder learning to disregard them. Therefore, when the AE training ends, the decoder is reset to its untrained state. This allows the decoder to learn from more meaningful latents provided by the encoder from the beginning.

#### S.1.2 Transformers

Transformer networks [19] operate on a set of elements, each represented by a vector of dimension *m*, which undergoes transformation by multiple transformer layers. To incorporate positional information and convert tokens into vectors, each token needs to undergo an embedding process, mapping it to a *m*-dimensional space using an embedding layer. The attention mechanism allows elements to selectively focus on and incorporate relevant information from other elements, facilitating effective information exchange and processing within the transformer architecture.

The two main categories of transformer networks were already outlined in the original transformer architecture [19], that described a translation system consisting of an encoder- and a decoder. The encoder evolved into the Bidirectional Encoder Representations for Transformers (BERT) architecture [32], while the decoder inspired the Generative Pretrained Transformer (GPT) family of auto-regressive models [33, 34, 35], including ChatGPT.

- **Encoder:** When translating from for example English to French, the encoder stack would take in the English text sequence and convert it into a sequence of contextual embeddings of the same length - only attending to all English tokens during the process.
- **Decoder:** In contrast, the decoder would in an auto-regressive way predicting each French word. This prediction is based on all previously generated French words as well as the whole encoding provided by the encoder for the English text. Therefore, two modifications to the attention mechanism are required:
- Firstly, the decoder is an AR model, meaning it predicts the next token in a sequence based on the preceding tokens. To train AR models efficiently, teacher forcing is commonly used. This means that the entire target sequence, like the French translation, is provided at once. To prevent the model from attending to future information during training, a masking mechanism is employed, setting the weights of future values to zero.
- Secondly, in translation tasks, the decoder needs to have access to the encoding of the original input sequence, such as the English sentence encoded by the encoder. This is achieved through cross-attention. This allows the decoder to attend to and incorporate information from a different sequence. In contrast, self-attention - which is also used in the encoder - refers to attending only within the same sequence.

In **Figure 8** we depict the details of the transformer blocks that are used to implement the encoder and decoder. Multiple sequential *Self Attention Blocks* form an *Encoder Stack* while multiple sequential *Cross Attention Blocks* form a *Decoder Stack*. Each transformer block updates a *Residual Stream* - a sequence of vectors received from the precedent transformer block or from an embedding layer. While the *Self Attention Block* (see **Figure 8a**) only considers information in the *Residual Stream* for updating the *Residual Stream*, the *Cross Attention Block* (see **Figure 8b**) also incorporates information from a separate stream (red lines in **Figure 8b**) into it.

**Figure 8:**
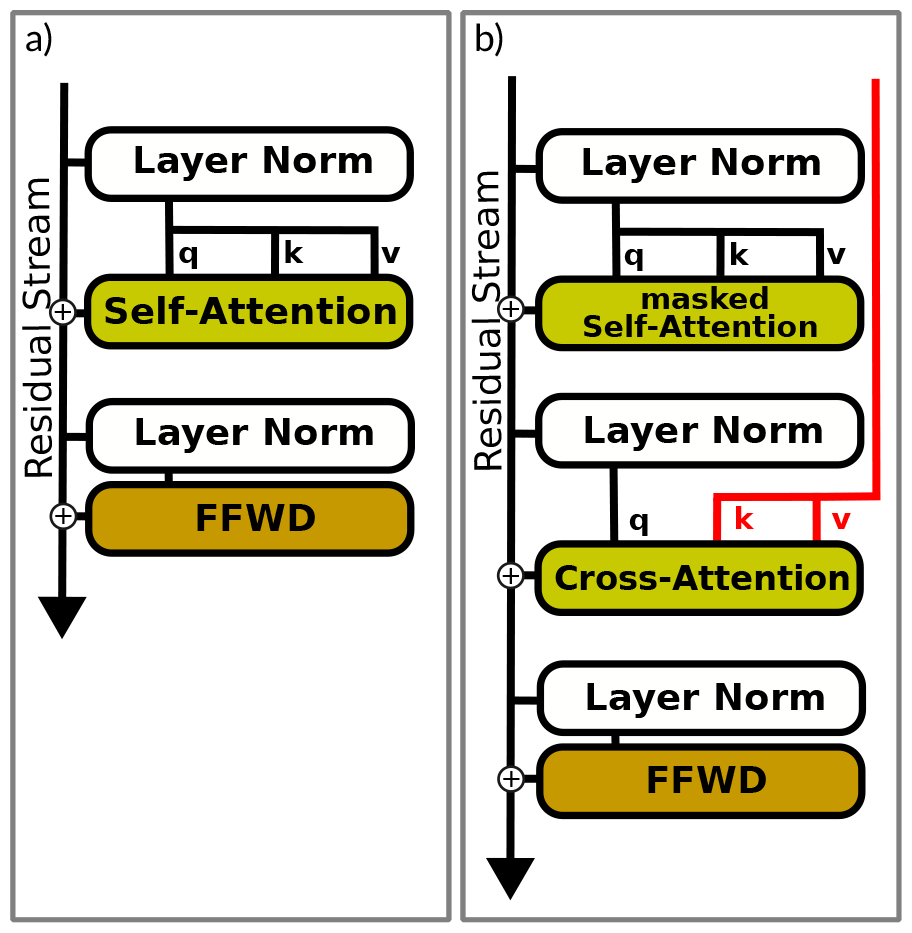
Transformer blocks add to a residual stream received from a preceding block or embedding layer. (a) The Self Attention Block uses self-attention only over the single input sequence itself. (b) In contrast, the Cross Attention Block applies attention two times. First, to self-attend over the residual stream sequence (this is typically done in a masked way - not attending over ‘future’ sequence elements that have not been generated yet). And second, to attend over a sequence that is different from the residual stream.

**Figure 9:**
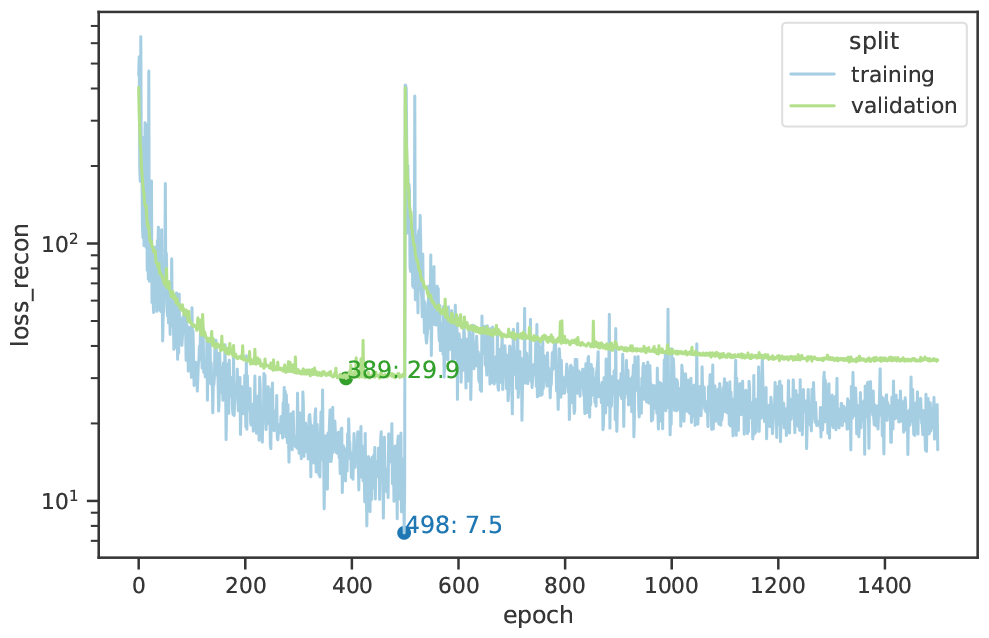
Reconstruction loss during training

#### S.1.3 Reinforcement Learning

We want to backpropagate gradient information from the MHC-I presentation predictor, through the `CAPE-XVAE` decoder, to the latent space. However, directly backpropagating through the discrete decoder output is not possible, so we require an alternative method. One such approach is RL, which involves learning how to take actions to maximize a reward signal [36]. RL has been utilized for training the model parameters of sequence-to-sequence models [37]. In contrast, we employ RL to train the latent representation *z*, which steers the decoders behavior.

The typical RL setting involves an agent observing its current state *s*_*t*_ and based on this taking an action *a*_*t*_ in an environment. The agent follows a policy *π*_*θ*_, parameterized by *θ*, to select an action, which can be stochastic. After performing the action, the environment provides the agent with a reward signal *r*_*t*_ based on the immediate benefit of the action in the environment. An episode consists of sequential iterations of this process until a terminal state is reached. The complexity of RL arises from the fact that actions taken at earlier stages of an episode can have repercussions on future rewards, either increasing or decreasing them. Consequently, the agent’s objective is to maximize the (sometimes discounted) expected sum of rewards (*J* (*θ*) = 𝔼 _*π*(*θ*)_ Σ_*t*_ *r*_*t*_) over the course of the episode. Additionally, we define the state value function *q*_*π*_(*s, a*), which represents the expected value of discounted rewards when taking action *a* in state *s* and subsequently following policy *π*. The state-value function serves as a useful estimate for evaluating the long-term desirability of actions within a given state.

The policy function *π*_*θ*_(*s*_*t*_) is crucial for maximizing the expected reward *J*. To find its optimal parameters *θ* (in our case the latent vector), the gradient ascent method shown in Equation 3 can be used:

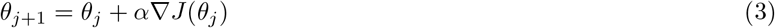

Directly back-propagating each trajectory through the discrete tokens is not possible. Fortunately, the REINFORCE [21] algorithm allows us to approximate ∇*J* (*θ*_*j*_) by the following Equation 4.

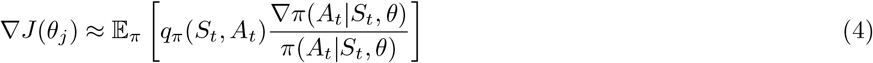

When we then define 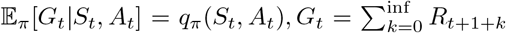 [36, Section 13.3], we get the following gradient ascent formula:

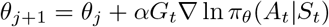

All of the terms therein can be calculated by us.

#### S.1.4 Training and hyper-parameter search

To optimize the model parameters we use Adam [38] (*β*_1_ = 0.9, *β*_2_ = 0.999) in combination with cosine annealing and utilize teacher-forcing during training.

To perform a systematic hyperparameter search, we employ Optuna [39] using the tree-structured parzen estimator method. The search objective is to minimize the VAE validation set loss including free bits.

In a first step we ran 51 trials of a hyper-parameter search across several dimensions to minimize the modified VAE loss on the validation set. We arrived at the following best hyper-parameter configuration in the column **Selected**.

Each of the trials was trained for 500 epochs on an auto-encoder objective (*β* = 0). Then, the decoder weights were reset to their initial state. Over the next 250 epochs of training the *β* parameter would linearly be increase to reach one. Afterwards training ran for another 750 epochs with *β* = 1. Figure 9 shows the development of the train and validation reconstruction losses during training. You can clearly see a spike when the `CAPE-XVAE`-decoder got reinitialized.

## Footnotes

1 https://services.healthtech.dtu.dk/services/NetMHCpan-4.1/

2 https://www.hiv.lanl.gov/components/sequence/HIV/search/search.html

3 https://new.rosettacommons.org/docs/latest/application_documentation/Application-Documentation

4 https:/seq2fun.dcmb.med.umich.edu/TM-align/

